# Oligomeric states in sodium ion–dependent regulation of cyanobacterial histidine kinase-2

**DOI:** 10.1101/140145

**Authors:** Iskander M. Ibrahim, Liang Wang, Sujith Puthiyaveetil, Norbert Krauß, Jon Nield, John F. Allen

## Abstract

Two-component signal transduction systems (TCSs) consist of sensor histidine kinases and response regulators. TCSs mediate adaptation to environmental changes in bacteria, plants, fungi, and protists. Histidine kinase 2 (Hik2) is a sensor histidine kinase found in all known cyanobacteria and as chloroplast sensor kinase in eukaryotic algae and plants. Sodium ions have been shown to inhibit the autophosphorylation activity of Hik2 that precedes phosphoryl transfer to response regulators, but the mechanism of inhibition has not been determined. We report on the mechanism of Hik2 activation and inactivation probed by chemical crosslinking and size exclusion chromatography together with direct visualisation of the kinase using negative-stain transmission electron microscopy of single particles. We show that the functional form of Hik2 is a higher order oligomer such as a hexamer or octamer. Increased NaCl concentration converts the active hexamer into an inactive tetramer. Furthermore, the action of NaCl appears to be confined to the Hik2 kinase domain.

**IMPORTANCE:** Bacteria sense change and respond to it by means of two-component regulatory systems. The sensor component is a protein that becomes covalently modified by a phosphate group on a histidine side chain. The response regulator accepts the phosphate group onto an aspartate, with structural and functional consequences, often for gene transcription. Histidine kinase 2 is a sensor of sodium ion concentration and redox potential, regulating transcription of genes for light-harvesting and reaction center proteins of photosynthesis in cyanobacteria and chloroplasts of algae and plants. Using radiolabeling, chemical crosslinking, chromatography and electron microscopy, we find that sodium ion concentration governs the oligomeric state of Histidine Kinase 2 and its phosphorylation by ATP.

## INTRODUCTION

Bacteria, algae, plants and fungi adapt to changes in their environments, and often utilise a sensor-response circuit known as a ‘two-component signal transduction system’ (TCS) to elicit physiological responses. TCSs are particularly diverse and widely distributed in bacteria. The simplest form of a TCS consists of just two proteins: a conserved sensor histidine kinase (component 1) and a response regulator (component 2).

A signal transduction cascade in bacteria usually begins at the cell membrane from where the signal propagates to the cytoplasm through the transmembrane domain of a sensor histidine kinase. The environmental cue is detected by the sensor domain located near the N-terminus of the histidine kinase polypeptide. Sensor histidine kinases also exist as soluble cytoplasmic proteins that perceive changes within the cell. Functional forms of both membrane-anchored and soluble histidine kinases occur predominantly as homodimers that contain conserved dimerization and phosphotransfer (DHp) and catalytic and ATP-binding (CA) domains. A higher-order oligomeric state, the tetramer, is promoted by an intermolecular disulfide bond in the histidine kinases DcuS (dicarboxylate uptake sensor and regulator) (1), RegB (Regulator B) (2), AtoS (Sensor kinase controlling ornithine decarboxylase antizyme) (3), and KdpD (Osmosensitive potassium channel sensor histidine kinase) (4). Formation of the higher-order oligomer appears to silence their autophosphorylation. In contrast, the well characterised soluble histidine kinases EnvZc (Core kinase domain of EnvZ) (5), VirAc (Virulence A) (6), and CheA (Chemotaxis histidine kinase A) (7) are reported to exist solely as inactive monomers or as active dimers. An *in vitro* study with the CheA sensor kinase indicates that its oligomeric state is concentration-dependent. CheA exists at low protein concentrations predominately as an inactive monomer while the extent of its dimerization increases with increasing protein concentration (7). Thus, CheA is likely to exist *in vivo* at equilibrium between its inactive monomeric and active dimeric forms, with its interaction with a ligand acting as a signal that shifts this equilibrium towards the active dimer. In contrast, the membrane-anchored sensor kinase DcuS from *E. coli* exists as monomer, dimer and tetramer both *in vitro* and *in vivo* (1). The ArcB sensor kinase of *E. coli* contains two conserved redox-active cysteines that are regulated by the redox state of ubiquinone. Oxidation of these cysteines leads to intermolecular disulfide bond formation between two monomers of ArcB, locking it into a tetrameric state inactive as a protein kinase (8, 9). The RegB histidine kinase of purple photosynthetic bacteria is also converted from an active dimer to an inactive tetramer by oxidation of its conserved cysteine (2).

Histidine kinase 2 (Hik2) is one of the three fully conserved histidine kinases found in cyanobacteria (10). The closest Hik2 homologue in nearly all algae and higher plants is Chloroplast Sensor Kinase (CSK) (11). A recombinant cyanobacterial Hik2 protein undergoes autophosphorylation on its conserved histidine residue and transfers the phosphoryl group to response regulators Rre1 and RppA (12). Rre1 is also modulated by Hik34 in response to increased temperature (13).

In its unmodified state, Hik2 appears to be autokinase active, and treatment with Na^+^ ions abolishes its autophosphorylation (12). The exact mechanism by which the activity of Hik2 is switched off by Na^+^ ions is not yet clear. Here we determine the mechanism of Hik2 activation and inactivation using chemical crosslinking and size exclusion chromatography, together with direct visualisation of the kinase using negative-stain transmission electron microscopy of single particles. We show that Hik2 is present in multiple oligomeric states *in vitro* and that a signal such as Na^+^ converts higher oligomers into the tetramer, thus inactivating it as the protein kinase that otherwise donates the phosphoryl group to its response regulators.

## RESULTS

### Distribution of Hik2 and domain architecture

Histidine kinases contain a conserved kinase core domain and a variable sensor domain. The kinase core domain is essential for autokinase and phosphotransfer activities. The Hik2 homolog that is present in almost all cyanobacteria and plants contains a conserved kinase core domain consisting of DHp and CA together with a GAF sensor domain (11, 12). In three cyanobacterial species Hik2 is present as a truncated form without a GAF sensor domain (14). We have examined the domain architecture and distribution of Hik2 homologs in cyanobacteria and chloroplasts. Hik2 domain architecture revealed that there are two forms of the Hik2 proteins in cyanobacteria and chloroplasts. Here these forms are designated class I and class II (Table 1 and Figure 1). Class I Hik2 proteins contain the full-length GAF domain as predicted with SMART database. Class II Hik2 proteins are predicted to have no GAF domain. As shown in Table 1, most cyanobacteria and chloroplasts have a class I Hik2. The least widely distributed Hik2 is the class II protein that is found only in three cyanobacteria, consistent with the finding of Ashby et al (14).

**Figure 1.**
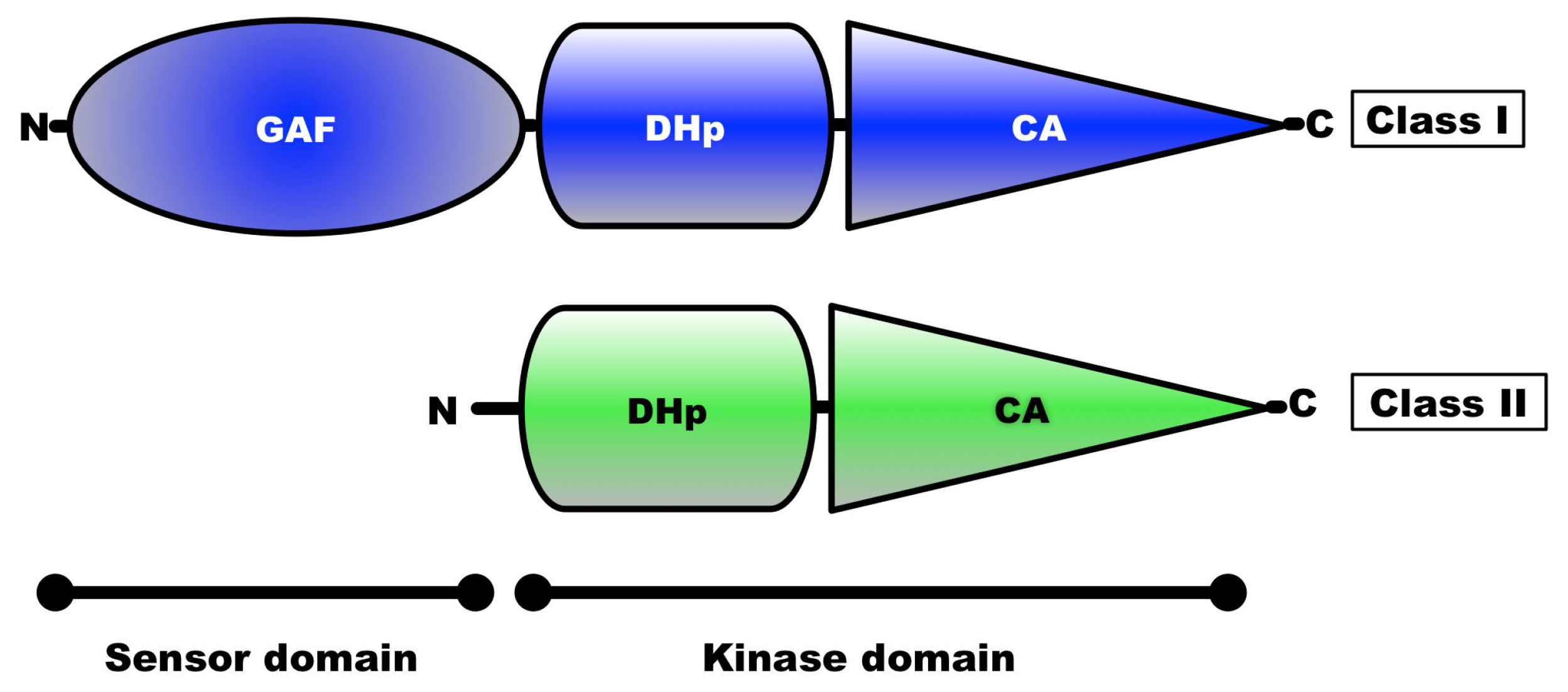
Domain Architecture of Hik2 Proteins. The amino and the carboxy termini are shown are shown as N and C, respectively. The domain architecture of Hik2 was predicted using the SMART database (33). The predicted sensor domain is shown as GAF. The kinase core contains the DHp and CA domains. The colour corresponds to different forms of Hik2 proteins; i.e. blue representing the full-length, class I Hik2 protein and green representing class II Hik2 protein.

**Table 1.** Distribution of full-length and truncated forms of Hik2. Tick (□) indicates the presence of Hik2.Full-length Hik2 proteins (forming Class I) contain the GAF domain as predicted with the SMART database. Truncated Hik2 proteins (forming Class II) are predicted to have no GAF domain. Names of chloroplast-containing species are coloured green.

| Species | Domain architecture of Hik2 |  |
| --- | --- | --- |
|  | Full-length | Truncated |
| *Synechocystis* sp. PCC 6803 | ☐ |  |
| *Synechococcus* sp. PCC 7002 | ☐ |  |
| *Cyanothece* sp. PCC7822 | ☐ |  |
| *Cyanothece* sp. PCC 8802 | ☐ |  |
| *Cyanothece* sp. PCC 8801 | ☐ |  |
| *Cyanothece* sp. PCC 7424 | ☐ |  |
| *Cyanothece* sp. ATCC 51142 | ☐ |  |
| *Cyanothece* sp. PCC 7425 | ☐ |  |
| *Crocospaera watsonii* WH0003 | ☐ |  |
| *Crocospaera watsonii*.WH8501 | ☐ |  |
| *Fischerella* sp. JSC-11 | ☐ |  |
| *Nostoc punctiforme* PCC 73102 | ☐ |  |
| *Nostoc* sp PCC 7120 | ☐ |  |
| *Cyanobacterium* UCYN-A | ☐ |  |
| *Cylindrospermopsis raciborskii*_CS-505 | ☐ |  |
| *Microcoleus chthonoplastes* PCC 7420 | ☐ |  |
| *Microcoleus vaginatus* FGP-2 | ☐ |  |
| *Microcystis aeruginosa* NIES-843 | ☐ |  |
| *Raphidiopsis brookii* D9 | ☐ |  |
| *Moorea product* 3L | ☐ |  |
| *Nodularia spumigena* CCY9414 | ☐ |  |
| *Anabaena variabilis* ATCC 29413 | ☐ |  |
| *Nostoc azollae* 0708 | ☐ |  |
| *Arthrospira maxima* CS-328 | ☐ |  |
| *Arthrospira maxima* CS-328 | ☐ |  |
| *Arthrospira* sp. PCC8005 | ☐ |  |
| *Arthrospira platensis* str. Paraca | ☐ |  |
| *Lyngbya* sp. PCC 8106 | ☐ |  |
| *Oscillatoria* sp. PCC 6506 | ☐ |  |
| *Thermosynechococcus elongatus* BP-1 | ☐ |  |
| *Trichodesmium erythraeum* IMS101 | ☐ |  |
| *Acaryochloris marina* MBIC 11017 | ☐ |  |
| *Synechococcus* sp. PCC7335 | ☐ |  |
| *Acaryochloris* sp. CCMEE 5410 | ☐ |  |
| *Synechococcus elongatus* PCC 7942 | ☐ |  |
| *Synechococcus elongatus* PCC 7942 | ☐ |  |
| *Synechococcus elongatus* PCC 6301 | ☐ |  |
| *Nodularia spumigena* CCY 9414 | ☐ |  |
| *Synechococcus* sp. WH 5701 | ☐ |  |
| *Synechococcus* sp. CB 0205 | ☐ |  |
| *Synechococcus* sp. RCC 307 | ☐ |  |
| *Synechococcus* sp. WH 8016 | ☐ |  |
| *Synechococcus* sp. WH7805 | ☐ |  |
| *Synechococcus* sp. CC9605 | ☐ |  |
| *Synechococcus* sp. WH 7803 | ☐ |  |
| *Prochlorococcus marinus* str. MIT 9301 | ☐ |  |
| *Prochlorococcus marinus* str. AS 9601 | ☐ |  |
| *Synechococcus* sp. JA-2-3B'a(2-13) |  | ☐ |
| *Synechococcus* sp. JA-3-3Ab |  | ☐ |
| *Gloeobacter violaceus* PCC 7421 |  | ☐ |
| *Cyanidioschyzon merolae* | ☐ |  |
| *Phaeodactylum tricornutum* | ☐ |  |
| *Thalassiosira pseudonana* | ☐ |  |
| *Oryza sativa* | ☐ |  |
| *Arabidopsis thaliana* | ☐ |  |
| *Populus trichocarpa* | ☐ |  |
| *Physcomitrella patens* | ☐ |  |
| *Ostreococcus tauri* | ☐ |  |
| *Ostreococcus lucimarinus* | ☐ |  |
| *Chlorella vulgaris* C-169 | ☐ |  |

### Recombinant protein production

The following proteins were cloned, overexpressed and purified from *E. coli*.: full-length *Synechocystis* sp. PCC 6803 (Syn_Hik2F); full-length *Thermosynechococcus elongatus* BP-1 Hik2 (Ther_Hik2F); the truncated core kinase domain of *T. elongatus* BP-1 protein containing only the DHp and CA subdomains (Ther_Hik2T); and the truncated DHp domain of *T. elongatus* BP-1 (Ther_DHp). Figure 2, lane 7 shows the purified Syn_Hik2F; lane 8, Ther_Hik2F; lane 9, Ther_Hik2T; lane 10 Ther_DHp. The calculated molecular weights are: Syn_Hik2F, 48.5 kDa; Ther_Hik2F, 44.2 kDa; Ther_Hik2T, 26.2 kDa; and Ther_DHp, 15.1 kDa. The apparent molecular weights on the SDS-PAGE are: Syn_Hik2F, 50 kDa; Ther_Hik2F, 45 kDa; Ther_Hik2T, 29 kDa; and Ther_DHp, 16 kDa.

**Figure 2.**
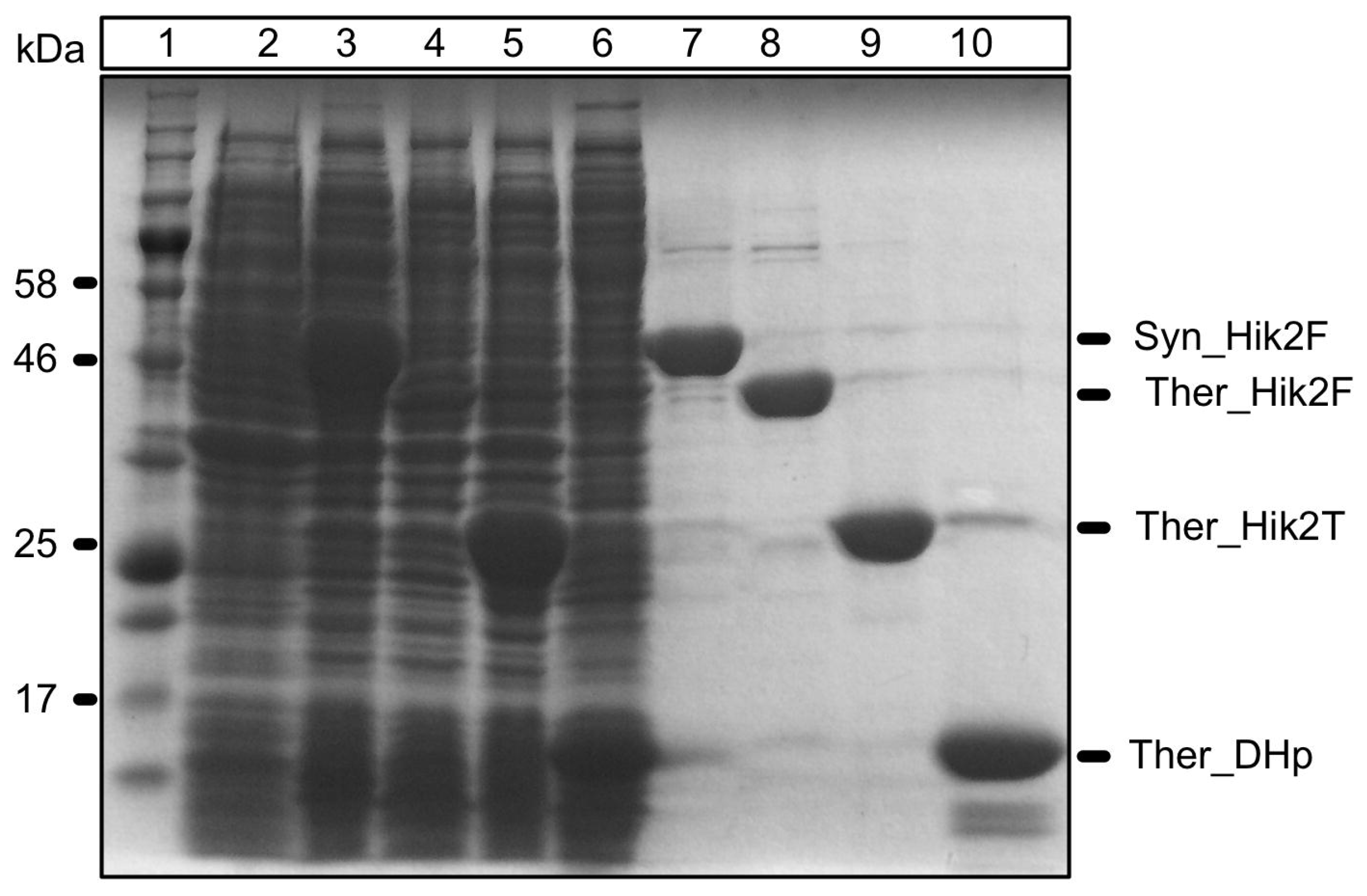
Protein overexpression and purification. The full-length *Synechocystis* sp. PCC 6803 and *Thermosynechococcus elongatus* BP-1 Hik2, and the truncated form of *Thermosynechococcus elongatus* BP-1 were overexpressed and purified as described in the experimental section. The following samples were loaded on a 10 % SDS-PAGE. Lane 1, protein molecular mass marker; lane 2 is total cell fraction before IPTG induction; lane 3–6 are total cell fractions after IPTG induction: lane 3, full-length *Synechocystis* Hik2 (Syn_Hik2F); lane 4, *Thermosynechococcus elongatus* BP-1 full-length Hik2 (Ther_Hik2F); lane 5, *Thermosynechococcus* truncated form (Ther_Hik2T); lane 6, *Thermosynechococcus* DHp domain (Ther_DHp); lane 7–10 correspond to elution fractions from the Ni^2^+ affinity chromatography column: lane 7, Syn_Hik2F; lane 8, Ther_Hik2F; lane 9, Ther_Hik2T; lane 10, Ther_DHp. The positions of the overexpressed proteins are indicated on the right, and the molecular masses of selected marker proteins are indicated on the left.

### Hik2 exists in higher-order oligomeric states

An earlier study has shown that sodium ions inhibit the autophosphorylation activity of full-length Hik2 (12). In order to elucidate its salt sensing mechanism, we investigated the oligomeric state of Hik2 in a buffer supplemented with or lacking NaCl. The oligomeric state of Hik2 was explored by size exclusion chromatography and chemical crosslinking with dithiobis(succinimidylpropionate) (DSP). These two techniques were chosen because size exclusion chromatography is most suited for stable protein-protein interactions, while DSP provides a more direct method of studying transient semi-stable protein-protein interactions. DSP is a symmetric molecule with two reactive groups connected by a spacer arm that is 12 Å in length. Thus DSP can form amide bonds with amino groups of two polypeptides that are in close proximity, for example, between two monomers in a dimer or between two dimers in a tetramer. DSP links polypeptides that interact under physiological conditions and therefore has advantages for studying oligomeric states of semi-stable protein-protein interactions.

In order to determine whether the oligomeric state of Hik2 is dependent on DSP concentration, Hik2 proteins were incubated with different concentrations of the crosslinker DSP for 10 minutes at 23 °C. Crosslinked products were then resolved on a non-reducing SDS-PAGE. Figure 3A, lane 2 shows that the untreated Hik2 protein migrated on a non-reducing SDS-PAGE with an apparent molecular weight of 50 kDa, corresponding to its monomeric form. Figure 3A, lanes 3–10 indicate that chemical crosslinking produced four distinct protein bands at apparent molecular weights corresponding approximately to multiples of 50 kDa. The first band at 50 kDa can be assigned to the monomer; a second band just above 190 kDa to a tetramer; and two further bands above 250 kDa to higher oligomers, possibly a hexameric and octameric form. Although increasing the concentration of DSP from 1 mM to 3 mM had no effect on the oligomeric state of Hik2 (Figure 3A, lanes 2–4), increasing the concentration above 3 mM resulted in a decrease in both monomeric and higher-order oligomers (Figure 3A, lane 3–12) and therefore only 2 mM of DSP was used in subsequent experiments.

**Figure 3.**
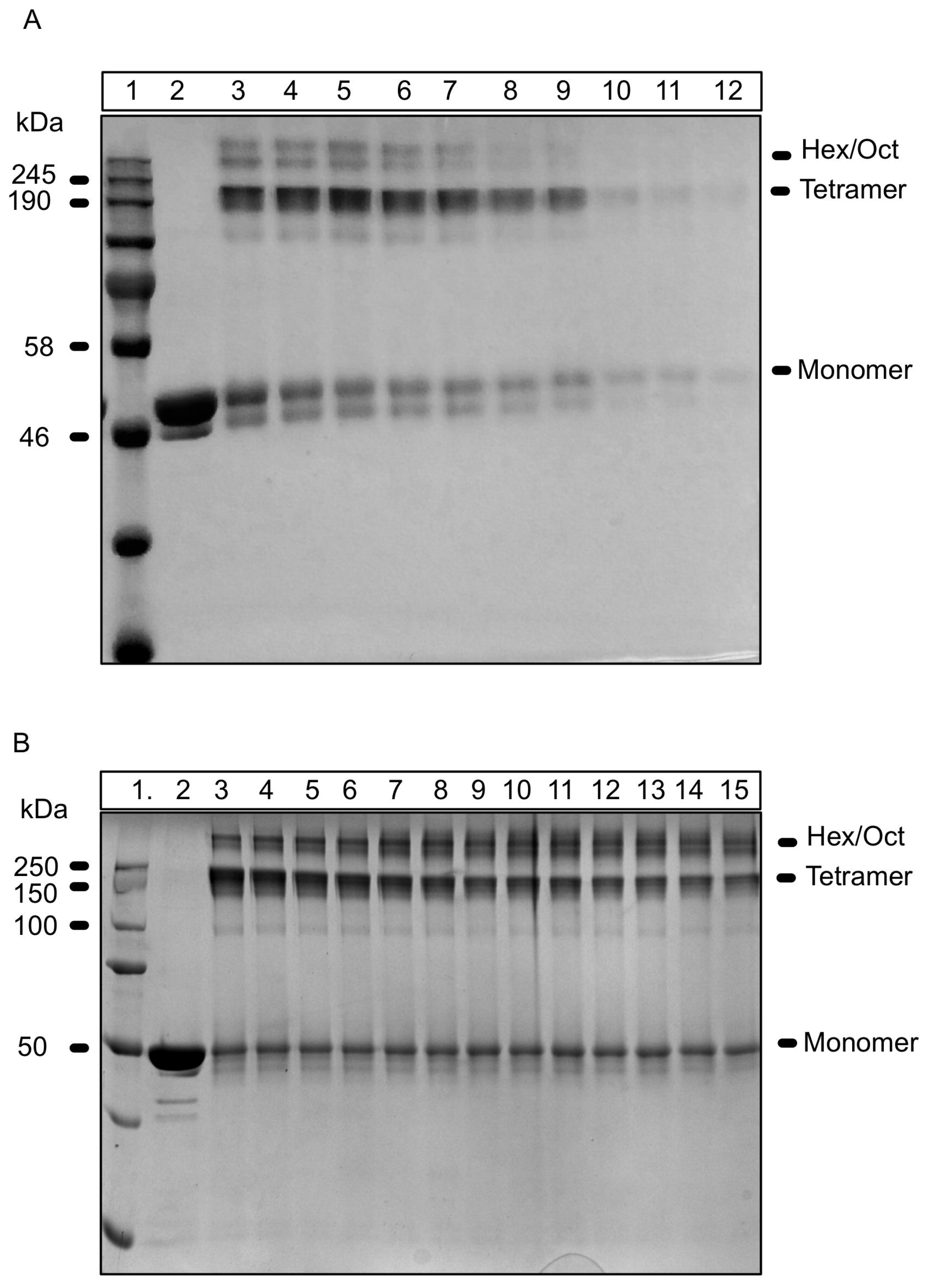
Effects of chemical crosslinking and protein concentration on the oligomeric state of *Synechocystis sp. PCC 6803* Hik2. A. Chemical crosslinking. Lane 1 shows protein molecular mass markers; lane 2, untreated Hik2 protein (control); lane 3, Hik2 treated with 1 mM DSP; lane 4, Hik2 treated with 2 mM DSP; lane 5, Hik2 treated with 3 mM DSP; lane 6, Hik2 treated with 4 mM DSP; lane 7, Hik2 treated with 5 mM DSP; lane 8 Hik2 treated with 6 mM DSP; lane 9, Hik2 treated with 7 mM DSP; lane 10, Hik2 treated with 9 mM DSP, lane 11, 10 mM treated with 10 mM DSP, lane 12 treated with 11 mM DSP. Samples were subjected to non-reducing 10 % SDS-PAGE. The molecular masses are shown on the left in kDa. The oligomeric states of Hik2 are indicated on the right. **B. Protein concentration**. Lane 1 shows protein molecular mass markers. Lane 2, 2 *μ*M untreated Hik2 protein. Proteins in the following lanes were cross-linked at the following protein concentrations: lane 3, 2 *μ*M; lane 4, 3 *μ*M; lane 5, 4 *μ*M; lane 6, 5 *μ*M; lane 7, 10 μM; lane 8, 15 *μ*M; lane 9, 20 *μ*M; lane 10, 25 *μ*M; lane 11, 30 μM; lane 12, 35 *μ*M; lane 13, 40 *μ*M; lane 14, 45 *μ*M; and lane 15, 50 μM. Molecular masses are shown on the left hand side in kDa. Different oligomeric states are labelled on the right hand side of the gel.

It has been shown that dimerization of CheA is concentration dependent (7). We therefore examined whether the oligomeric state of Hik2 depends on its concentration. Crosslinking was performed with differing concentrations of Hik2 proteins, ranging from 2 to 50 *μ* M, while the concentration of DSP, incubation times, and temperature were kept constant. Equal quantities of crosslinked Hik2 proteins were then analysed with non-reducing SDS-PAGE. No correlation is observed between the apparent quantity of the monomeric state of Hik2 and protein concentration while the quantity of the tetramer appears to decrease and hexamer and octamer increased with increasing protein concentration (Figure 3B).

### Monomer, tetramer, and hexamer forms of Hik2 are autokinase active and salt converts the higher-order oligomers into a tetramer

In order to investigate the functional states of Hik2 oligomers, we carried out autophosphorylation of Hik2 before and after it was crosslinked. Figure 4A, lane 2, shows Hik2 protein that was allowed to autophosphorylate and then crosslinked with DSP. The result was phosphorylated monomeric, tetrameric, and higher order oligomeric forms. Figure 4A, lane 3, shows that Hik2 protein, which was first crosslinked and then subjected to the autophosphorylation assay, produced monomers, tetramers and higher oligomers relatively inactive in autophosphorylation. Thus crosslinking may lock the protein into an inactive state. Since the activity of Hik2 is suppressed by salt (12), we then investigated the effect of salt on the oligomeric state of Hik2. Figure 4B, lane 2, shows NaCl untreated and non-crosslinked Hik2 protein migrating as monomers, and in lane 3 NaCl untreated but crosslinked protein migrating as monomers, tetramers and higher-order oligomers as expected. Lane 4 shows Hik2 that was treated with NaCl first and then crosslinked. Addition of NaCl appears to have resulted in the conversion of the hexamers and octamers into tetramers. The proportion of the monomeric form is also decreased in figure 4B, lane 4 in the salt-treated sample.

**Figure 4.**
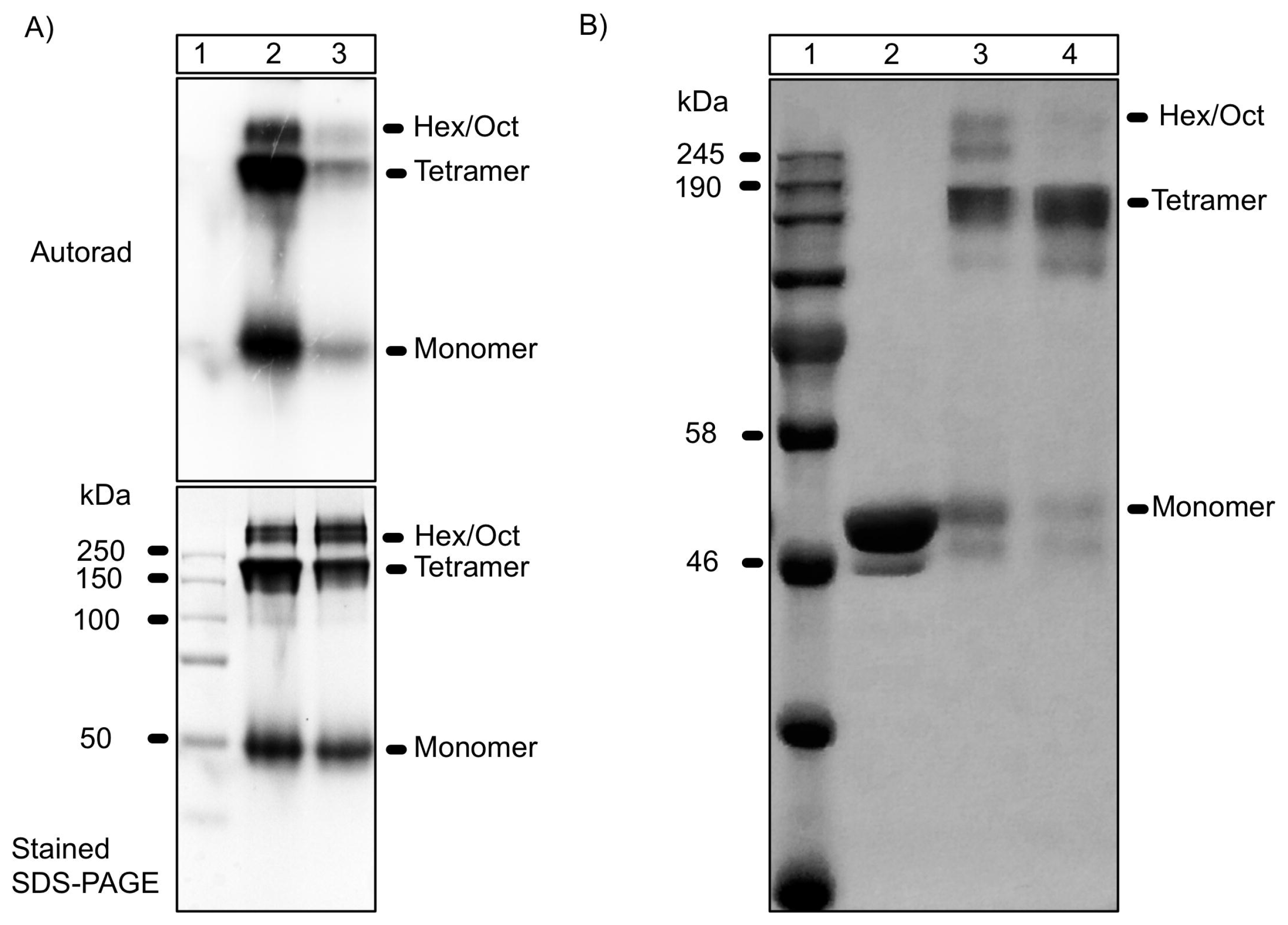
Functional characterisation of oligomeric states of *Synecftocystis* sp. PCC 6803 Hik2. A) *Autophosphorylation activity of Hik2*. Lane 1 shows protein molecular mass markers; lane 2, Hik2 protein that was allowed to autophosphorylate before crosslinking; lane 3, Hik2 protein that was first crosslinked, followed by autophosphorylation. B) *Effect of salt on the oligomeric state of Hik2*. Lane 1 shows protein molecular mass markers; lane 2, shows untreated Hik2 protein (control); lane 3, salt untreated Hik2 crosslinked; lane 4, salt treated Hik2 crosslinked. Different oligomeric states are labelled on the right hand side of the gel.

### NaCl converts the active hexamer form of Hik2 into a tetramer

On a Superdex 200 column calibrated with buffer lacking NaCl the full-length *Synechocystis* and *Thermosynechococcus* and the truncated *Thermosynechococcus* Hik2 proteins eluted as hexamers with apparent molecular weights of 380 kDa, 260 kDa and 150 kDa, respectively (Figure 5A, 5B and 5C, solid lines). In the presence of NaCl, the apparent molecular weight was shifted from 380 to 200 kDa for *Synechocystis* Hik2, from 260 to 200 for *Thermosynechococcus* Hik2, and from 150 to 100 for the truncated *Thermosynechococcus* Hik2, corresponding to tetrameric forms (Figure 5A, 5B and 5C, broken lines). These results may be consistent with those obtained from crosslinking experiments (Figure 4B), provided that one assumes that contributions from tetramers to overlapping bands are difficult to resolve in the samples untreated with NaCl. Since the DHp domain of the histidine kinase is important for its dimerisation activity, we explored the role of the DHp domain in higher order oligomeric formation. Figure 5D shows that the DHp domain of *Thermosynechococcus* formed high order oligomers that may be octamers in buffer without NaCl, and that the ‘octamers’ converted to ‘hexamers’ upon treatment with NaCl. It may be concluded that the DHp domain is of central importance for the salt sensing activity of Hik2. The oligomeric states observed for the DHp domain are different from those seen for the full-length proteins, and it is possible that these are controlled by additional interactions involving other domains. It was noted that oligomeric states equivalent to the full-length proteins were only observed for the truncated *Thermosynechococcus* Hik2 (Thermo_Hik2T) form, thus it is possible the CA domain may play a role in defining the Hik2 oligomeric state.

**Figure 5.**
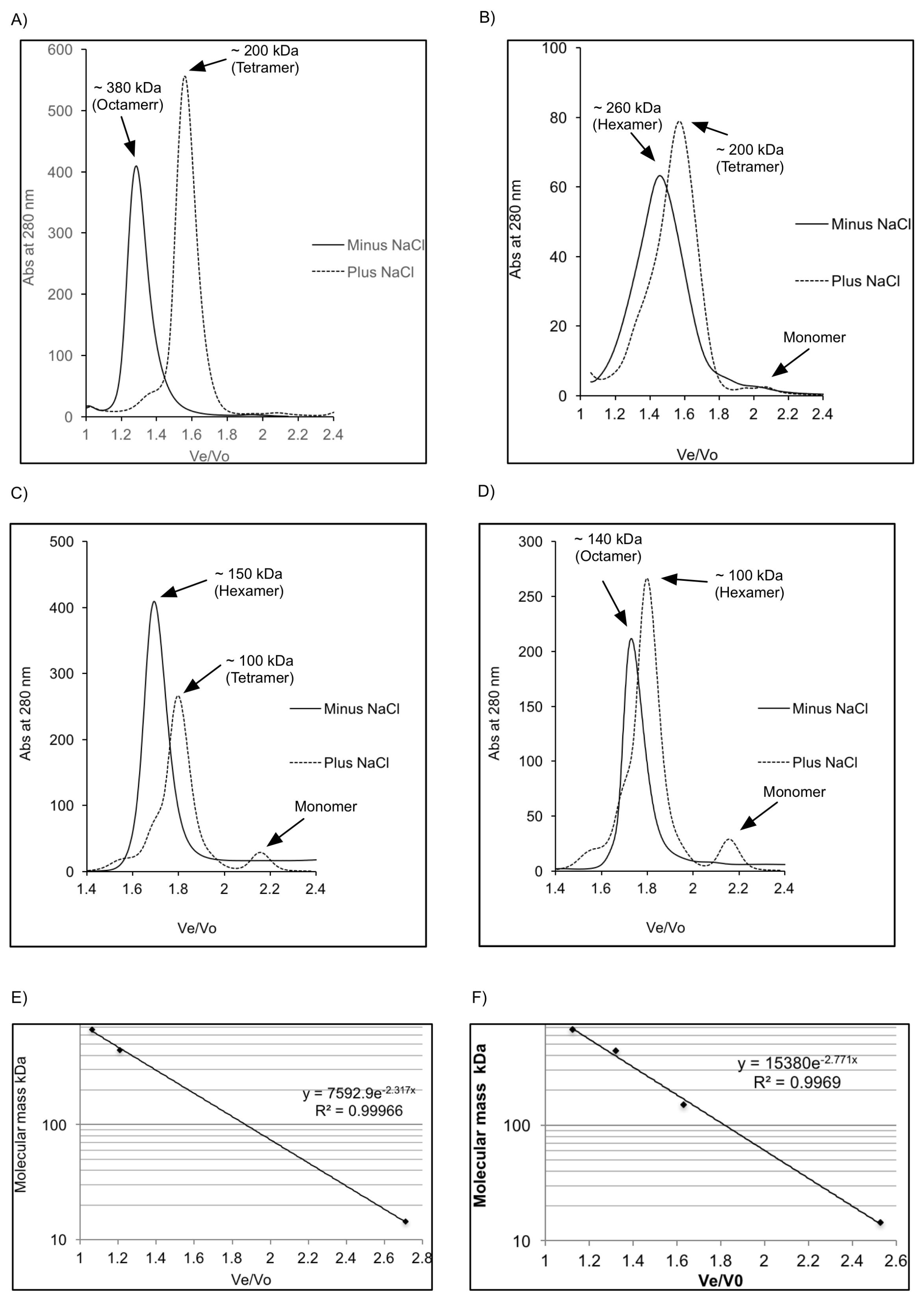
Separation of Hik2 oligomers by size exclusion chromatography. Typical elution profiles of Hik2 proteins on a Superdex 200 column eluted with a buffer containing 20 mM Tris-HCl (pH 7.6) and 10 mM NaCl (solid line) or with 20 mM Tris-HCl (pH 7.6) and 500 mM NaCl (dotted line). **A**) Syn_Hik2F; **B**) Ther_Hik2F; **C**) Ther_Hik2T; **D**) Ther_DHp. The positions of hexamers, tetramers and octamers are shown. **E**) Calibration curve (E, 10 mM NaCl, and F, 500 mM NaCl) of the Superdex 200 using standard proteins of known molecular weight: Apoferritin (443 kDa), Alcohol dehydrogenase (150 kDa), and Carbonic anhydrase (29 kDa). Blue dextran (2000 kDa) was used to determine the void volume (Vo). Ve is the effluent volume. On the y-axis the base ten logarithm of the protein molecular mass is shown, and on the x-axis Ve/Vo is shown.

### Transmission electron microscopy (TEM) and single particle analysis of negatively stained Hik2

Five independent samples were negatively stained and imaged using a JEOL 1230 TEM equipped with a 2k Olympus Morada CCD camera system and those micrographs that displayed the highest quality (41 from a total of 201, see Methods) were carried forward for single particle image analysis. Figure 6 is a micrograph of a negatively stained *Synechocystis* Hik2 protein sample as observed by TEM at 80,000 × magnification and typical for those used in the single particle image-averaging analysis. The high contrast produced by the uranyl acetate stain allowed for the visual inspection of protein complexes. A variety of different sizes and orientations were observed. A dataset of 13,341 individual protein complex images was built and subjected to reference-free alignment and multivariate statistical classification using Imagic-5 software. The spread of oligomeric states, i.e. structural heterogeneity, may be appreciated so that the TEM-derived dataset is represented as 600 class averages after four rounds of iterative refinement (Figure 7) After four rounds of iterative refinement, unassigned particles were excluded leaving 11,371 particles that were classified into 100 class averages, or ‘characteristic views’ (Figure 8 and Table 2). Each row of Figure 9 depicts four of these characteristic views for each of the different oligomeric complex families assigned subjectively; monomers, dimers, trimers, double dimers or hexamers, and broken or ambiguous density, respectively. It cannot be ruled out that the oligomers shown in Figures 9B and 9C are double dimers (dimers of dimers) and double trimers (trimers of dimers), respectively, viewed in projection. Similarly, it is possible that the oligomers shown in Figure 9C are double trimers or double tetramers (dimers of tetramers) viewed in side elevation.

**Figure 6.**
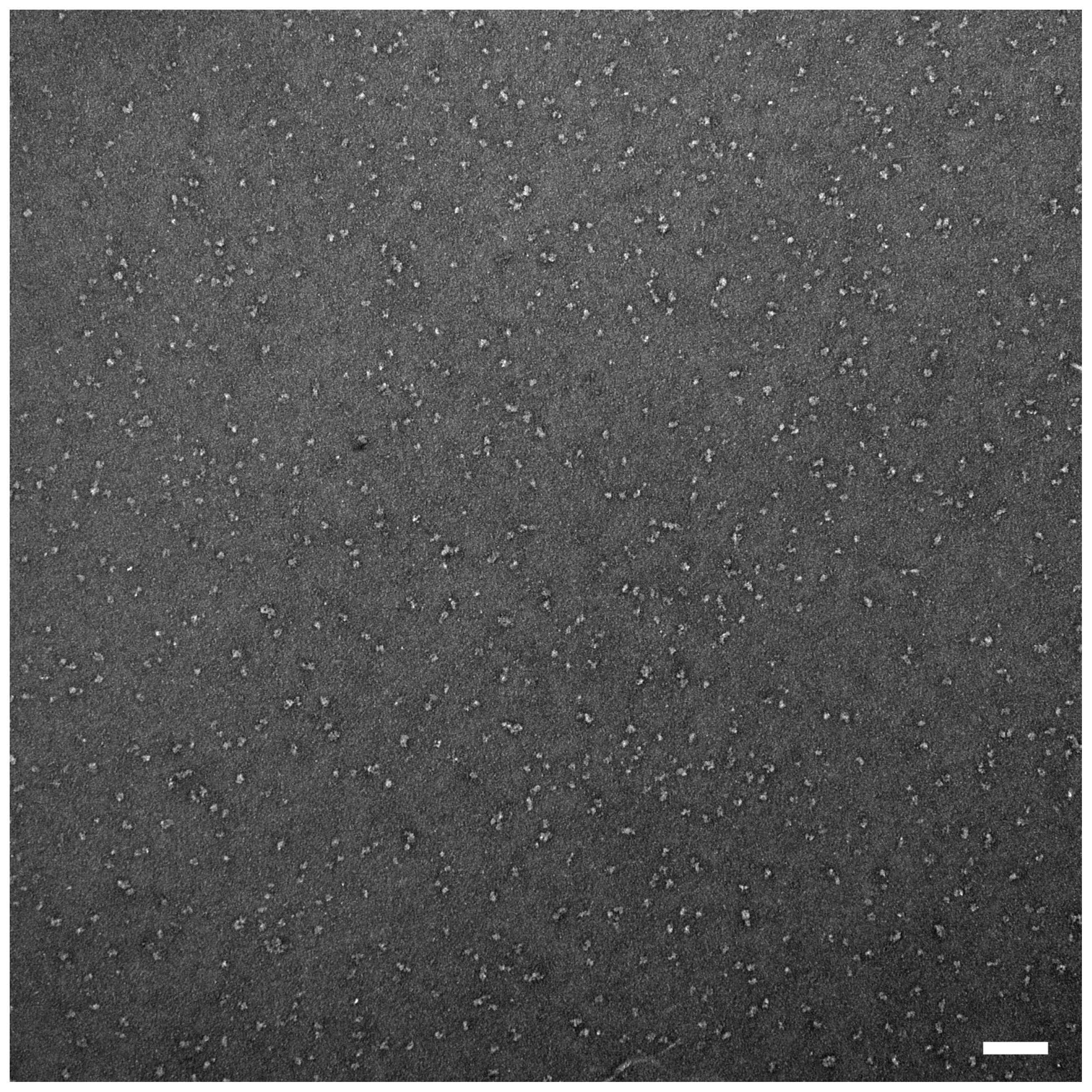
Characteristic micrograph of a negatively stained *Synechocystis* Hik2 protein sample. A typical micrograph as observed by TEM at 80,000 × magnification. Bar represents 100 nm.

**Figure 7.**
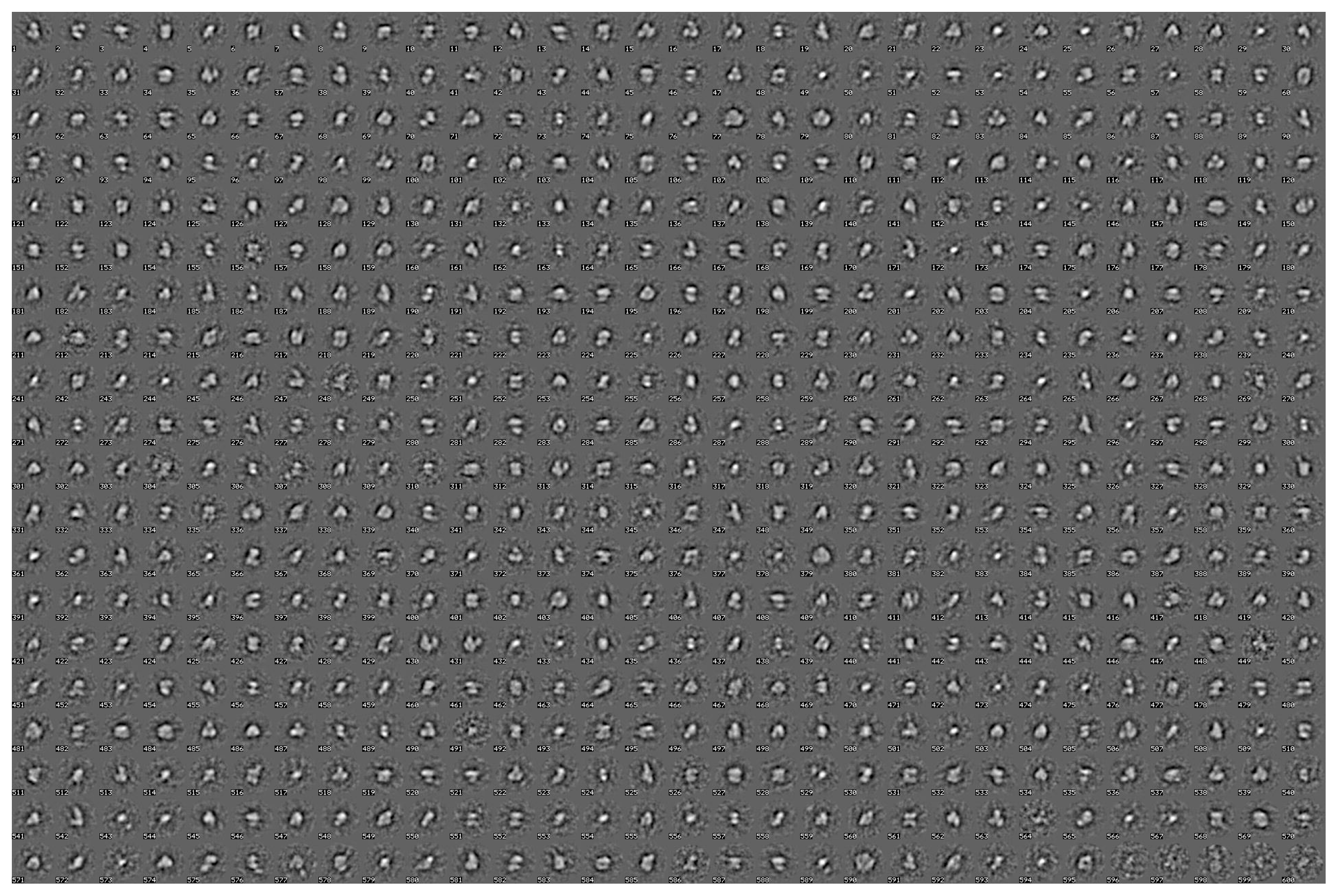
Structural heterogeneity of the Hik2 single particle TEM-derived dataset. The structural heterogeneity was revealed by relaxing the classification constraints, presenting all possible single particles automatically particle-picked from the micrographs as 600 characteristic views (class averages). Each boxed side of a single characteristic view represents 382 Å.

**Figure 8.**
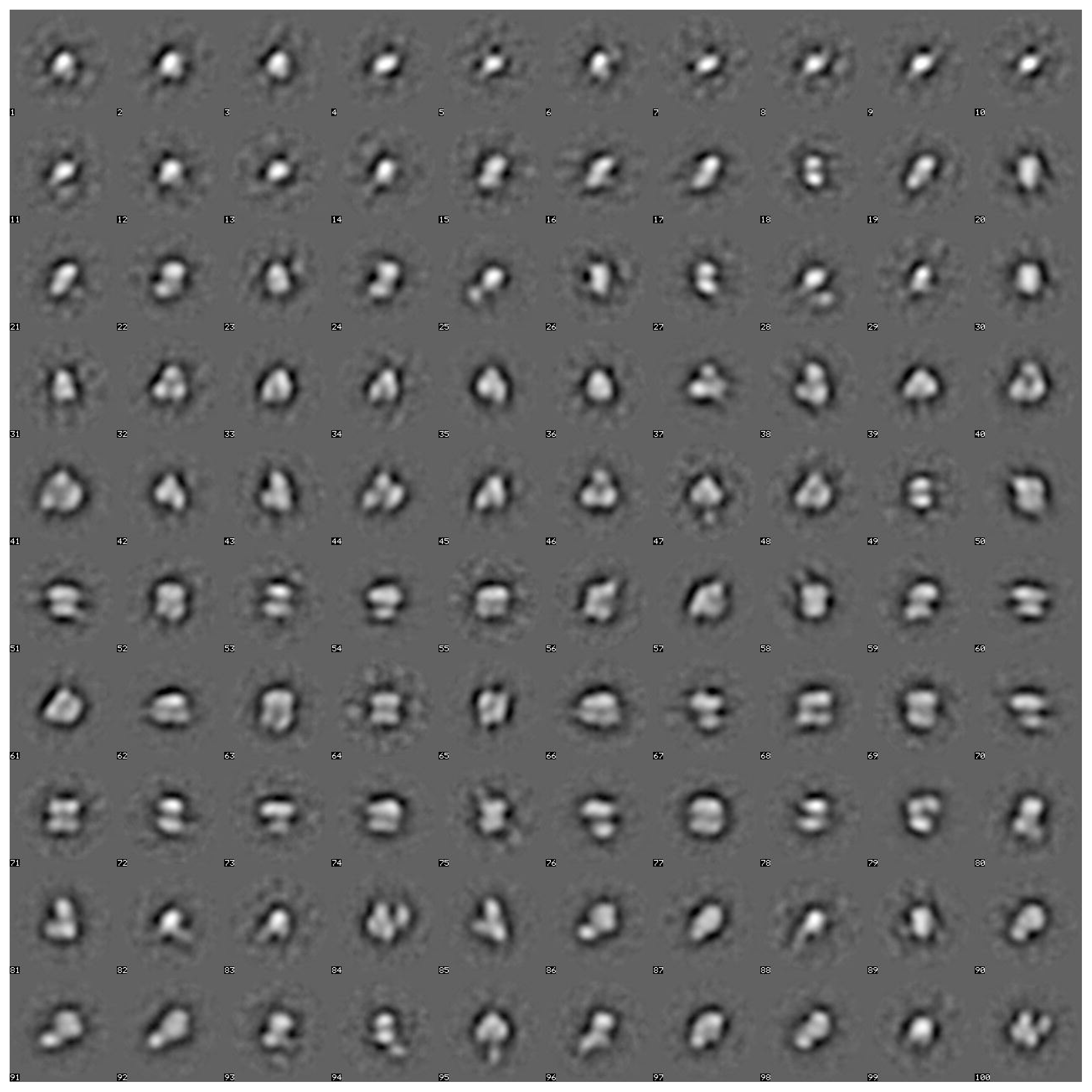
Classification of the Hik2 single particle TEM-derived dataset. After 4 rounds of iterative refinements, the dataset was classified into 100 characteristic views (class averages). These views were re-ordered by visual inspection into oligomers of increasing diameter. Statistical analysis of this classification (see Table 2) revealed that these views represent 11,371 particles, which were subsequently attributed to oligomers ranging from monomers through to double-trimers (hexamers), and stacked/double tetramers (octamers). The side of each box, within which individual characteristic views are floated, represents 382 Å in length, thus a typical monomer is ~80 Å in diameter and the largest views are ~180 Å in long axis.

**Figure 9.**
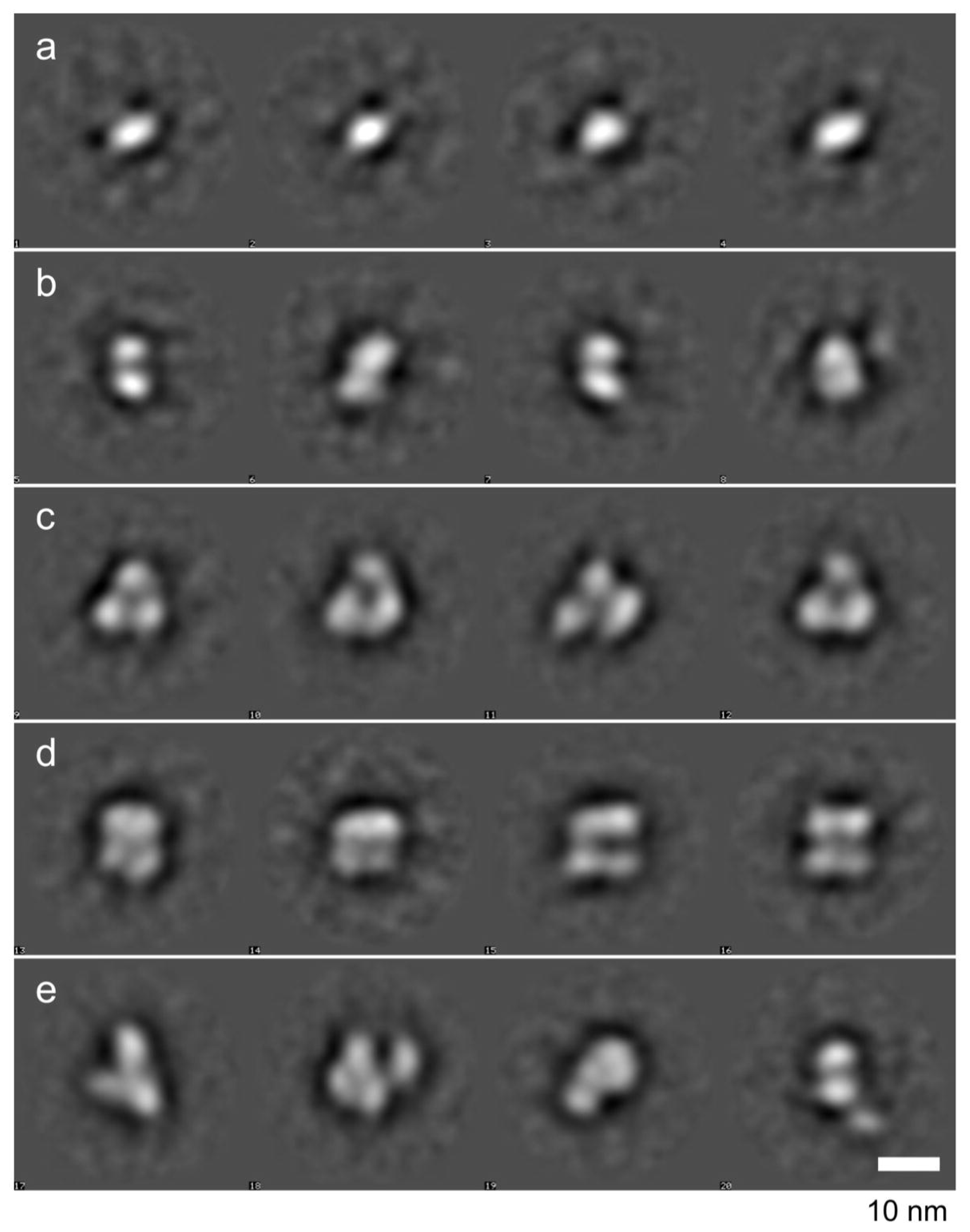
Single particle averages of negatively stained samples imaged by transmission electron microscopy (TEM). After the image-processing techniques of multivariate statistical analysis and subsequent averaging a series of Hik2 protein complex subpopulations or families was observed by TEM of negatively stained samples. Comparison of this figure with Table 2 provides a detailed statistical outcome for each subpopulation. Four representative averages for each family are shown. These are attributed to oligomeric states of a) monomers, b) dimers, c) trimers or potential double trimers (hexamers) in projection, d) tetramers, potential double trimers (hexamers) or double tetramers (octamers) in side elevation, e) characteristic views of the remaining family of broken complexes or those with extraneous density visible. The *Synechocystis* sp. PCC 6803 Hik2 monomer has a molecular mass of ~50 kDa. Protein is white, stain black. The scale bar is 10 nm.

**Figure 10.**
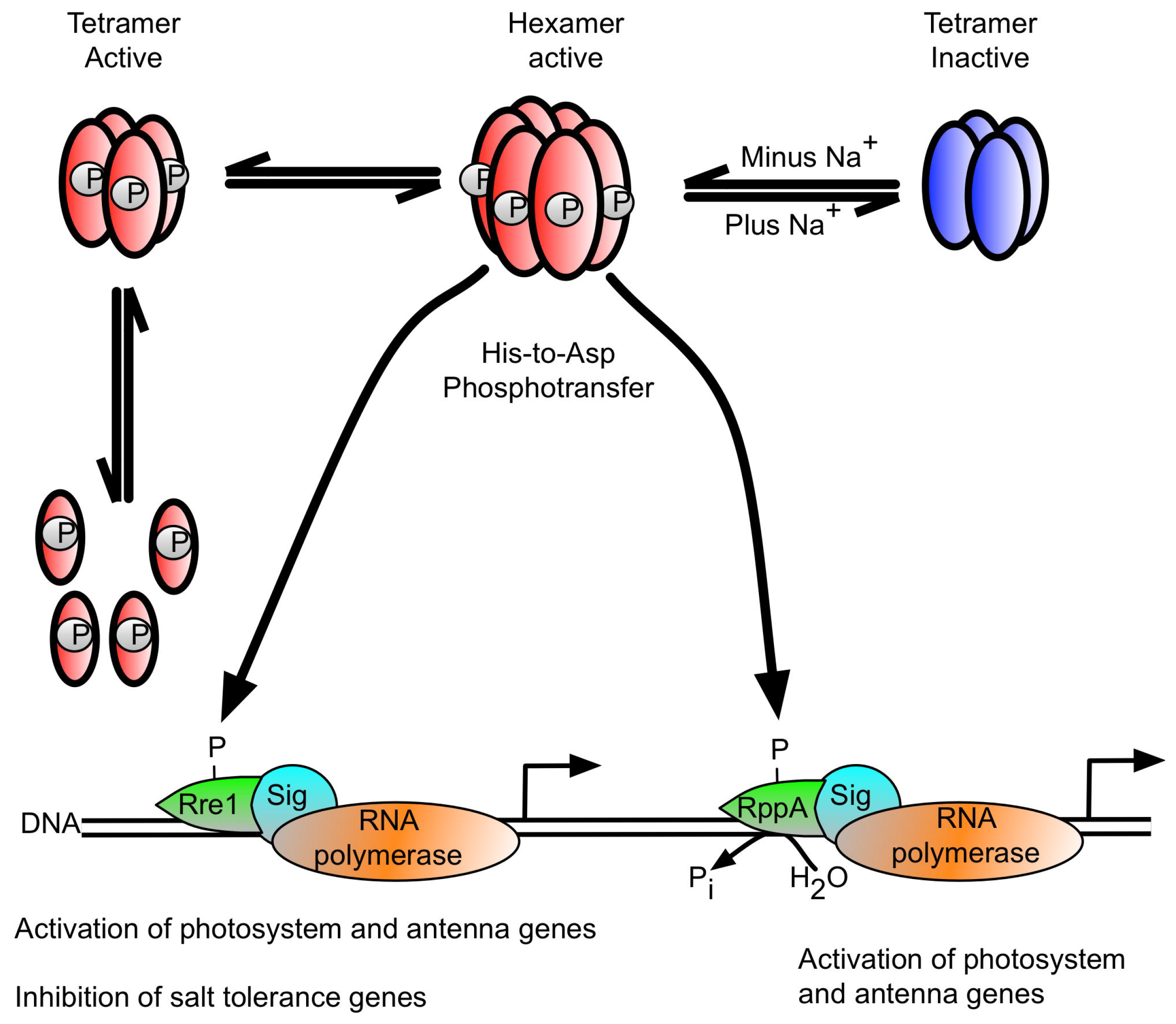
Proposal for a Hik2-based signal transduction pathway in cyanobacteria. The hexameric form of Hik2 is autokinase active and the oligomeric state of Hik2 is regulated by signals from Na^+^ and the photosynthetic electron transport chain. The active hexameric form of Hik2 is converted to an inactive tetramer upon salt stress.

**Table 2.**
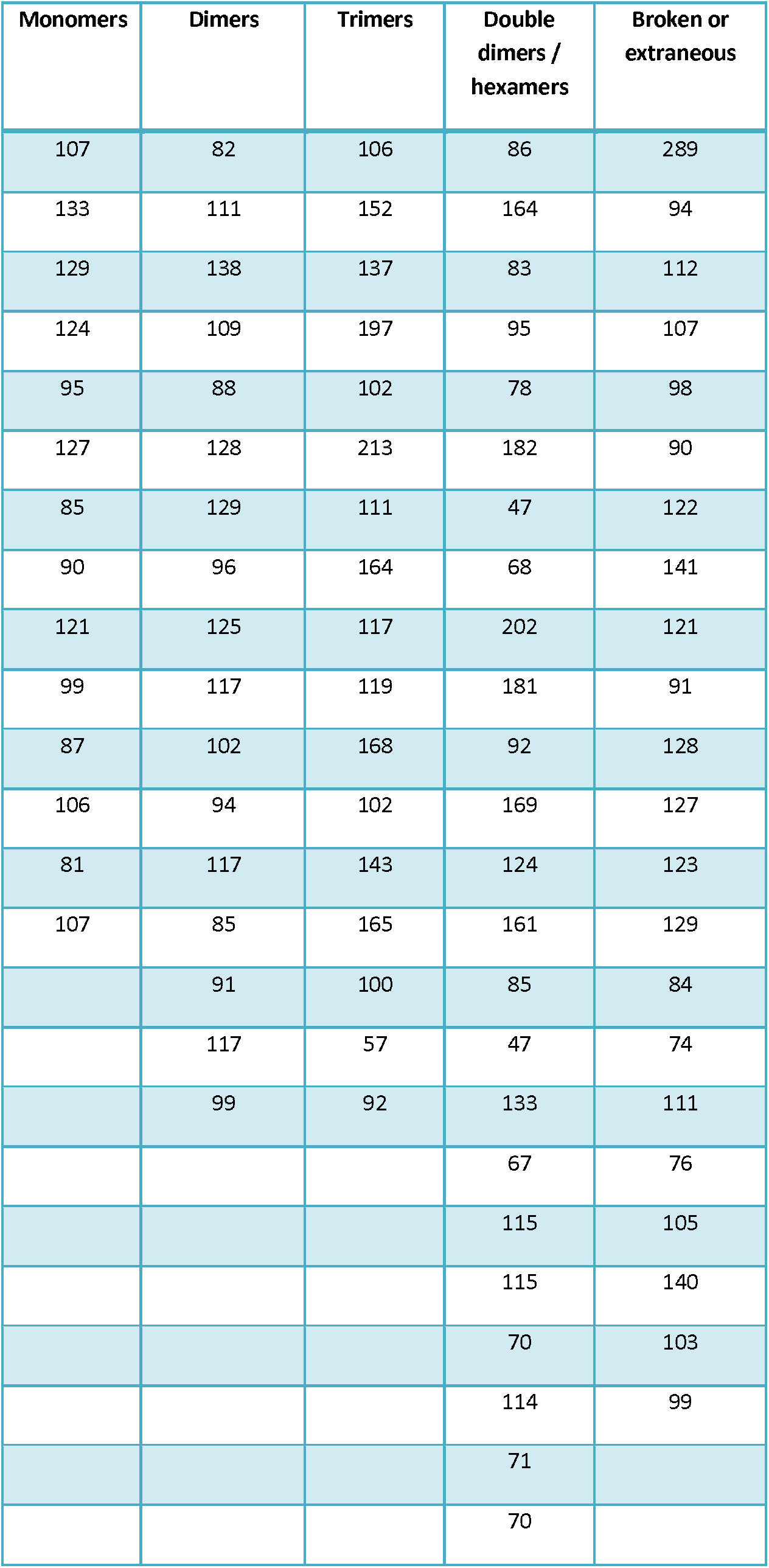
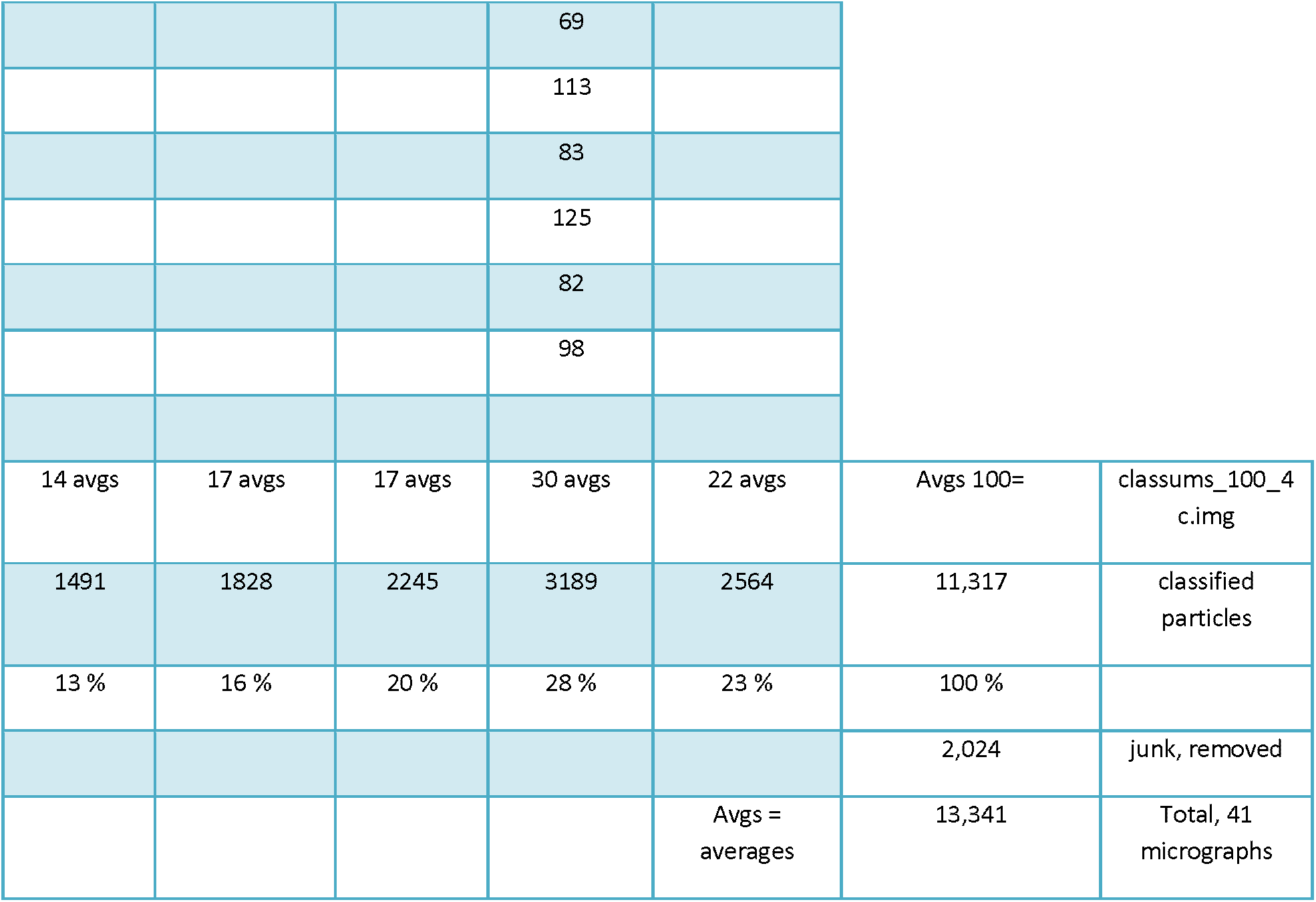
Single particle image processing statistics relating to the characteristic views (averages) presented in Figure 8 (100 averages), the final classification of the dataset derived from micrographs of which Figure 6 is typical. Each number below refers to the single particles present in each average; e.g. the first monomer average, contains 107 particles.

| Monomers | Dimers | Trimers | Double dimers / hexamers | Broken or extraneous |
| --- | --- | --- | --- | --- |
| 107 | 82 | 106 | 86 | 289 |
| 133 | 111 | 152 | 164 | 94 |
| 129 | 138 | 137 | 83 | 112 |
| 124 | 109 | 197 | 95 | 107 |
| 95 | 88 | 102 | 78 | 98 |
| 127 | 128 | 213 | 182 | 90 |
| 85 | 129 | 111 | 47 | 122 |
| 90 | 96 | 164 | 68 | 141 |
| 121 | 125 | 117 | 202 | 121 |
| 99 | 117 | 119 | 181 | 91 |
| 87 | 102 | 168 | 92 | 128 |
| 106 | 94 | 102 | 169 | 127 |
| 81 | 117 | 143 | 124 | 123 |
| 107 | 85 | 165 | 161 | 129 |
|  | 91 | 100 | 85 | 84 |
|  | 117 | 57 | 47 | 74 |
|  | 99 | 92 | 133 | 111 |
|  |  |  | 67 | 76 |
|  |  |  | 115 | 105 |
|  |  |  | 115 | 140 |
|  |  |  | 70 | 103 |
|  |  |  | 114 | 99 |
|  |  |  | 71 |  |
|  |  |  | 70 |  |

**Table 2.** Single particle image processing statistics relating to the characteristic views (averages) presented in Figure 8 (100 averages), the final classification of the dataset derived from micrographs of which Figure 6 is typical.
|  |  |  |  |  |  |  |
| --- | --- | --- | --- | --- | --- | --- |
|  |  |  | 69 |  |  |  |
|  |  |  | 113 |  |  |  |
|  |  |  | 83 |  |  |  |
|  |  |  | 125 |  |  |  |
|  |  |  | 82 |  |  |  |
|  |  |  | 98 |  |  |  |
| 14 avgs | 17 avgs | 17 avgs | 30 avgs | 22 avgs | Avg 100= | classums_100_4<br>c.img |
| 1491 | 1828 | 2245 | 3189 | 2564 | 11,317 | classified<br>particles |
| 13 % | 16 % | 20 % | 28 % | 23 % | 100 % |  |
|  |  |  |  |  | 2,024 | junk, removed |
|  |  |  |  | Avg =<br>averages | 13,341 | Total, 41<br>micrographs |

## DISCUSSION

It has been suggested that membrane bound and soluble histidine kinases are homodimeric in their functional forms (15–17) and that some histidine kinases interconvert between an inactive monomer and an active dimer (7). Although histidine kinases are also reported to exist in higher order oligomeric states, these states seem to be autokinase inactive. The exception is the hybrid histidine kinase ExsG, which has been shown to be active as a hexamer (18). The work presented here was directed at elucidation of the oligomeric states of the soluble cyanobacterial Hik2 protein. Using chemical crosslinking, size exclusion chromatography profiles, and transmission electron microscopy, we find that the full-length Hik2 protein exists as tetramers, hexamers and other higher-order oligomers (Figures 3, 5 and 7). Further analysis of the oligomeric states of Hik2 using truncated forms of the protein revealed that the oligomeric state of Hik2 is controlled by its kinase domain (Figures 5C and 5D), and that treatment with NaCl converts the hexamer into a tetramer (Figures 4B and 5), rendering the kinase inactive. Furthermore, the Hik2 protein is not always present in its full length form, and may be truncated in certain photosynthetic organisms (Table 1).

The occurrence of Hik2 in different forms implies that it may have multiple functions, some specific to certain species. The class I Hik2 protein homologs, which are found in chloroplasts and cyanobacteria, have a fully conserved GAF domain as their sensor domain and may therefore be able to sense a variety of signals. The class II Hik2, found only in marine cyanobacteria, have lost the GAF sensor domain completely (Table 1 and Figure 1). Based on the results in Figures 5C and 5D, the activity of this class II Hik2 may be regulated through its kinase domain. Class II Hik2 are found only in three cyanobacterial species; *Gloeobacter violaceus* PCC 7421, *Synechococcus* sp. JA-2-3B’a(2–13), and *Synechococcus* sp. JA-3-3Ab. Current phylogenetic trees indicate that these three cyanobacteria diverged very early from other cyanobacteria (19). It is therefore possible that the ancestral Hik2 protein lacked a sensor domain and later acquired the GAF sensor domain through gene fusion or acquisition. Alternatively, the sensor domain of class II Hik2 within those three cyanobacteria might have been lost after they diverged from a common ancestor with a sensor domain-containing Hik2.

The activity of histidine kinases is modulated by environmental cues through signal-induced conformational changes (20–22). Chemical crosslinking and size exclusion chromatography were utilised in order to understand the effect of salt stress on the oligomeric state an activity of Hik2. Both techniques revealed that the Hik2 protein complex exists predominantly as tetrameric, hexameric and other higher order oligomeric forms (Figures 3 and 5). Monomers are visible in Figure 3A and 3B, lane 3. However, salt treatment decreased the monomeric forms of Hik2 (Figure 4B, lane 4). Interestingly, unlike the histidine kinases CheA (7), DcuS (1), ArcB (8, 9), and RegB (2) the dimeric form of Hik2 was not detected, indicating that higher order oligomers are the stable forms. Furthermore, the higher order oligomeric states of Hik2 were functionally active (Figure 4A). Treatment of Hik2 with NaCl converted the monomer, hexamer and octamer into the tetramer (Figure 4B, lane 4, and Figure 5). Results obtained from crosslinking (Figure 4) and size exclusion chromatography (Figure 5) indicate that the inactivation mechanism of Hik2 involves the conversion of the higher order oligomers into the tetramer. What remains to be clarified, however, is how it is possible that tetramers in Hik2 samples that were not treated with NaCl exhibit autokinase activity, suggesting that these tetramers are structurally different from the inactive tetramers that form at elevated NaCl concentrations.

The sensor domain of histidine kinases is usually required to detect signals, the exception being EnvZ, which was shown to receive signal through its DHp domain (23). It is therefore likely that Hik2 employs a DHp-based signal perception mechanism for its salt sensing activity. We tested this possibility using three different variants of the Hik2 protein. Our result showed that the truncated forms of Hik2, consisting of the core kinase domain or DHp domain alone, were both present as a higher order oligomer and that treatment with NaCl converted them into the lower form of oligomer. We conclude that that the salt sensing activity of Hik2 is confined to its DHp domain. In addition to salt, the GAF domain of Hik2, which is found in class I Hik2s, might be involved in perceiving additional signal(s) such as redox or it might bind small ligand(s) required to regulate its autophosphorylation (24). Indeed, the Hik2 homologue in higher plants has been shown to bind the PQ analogue DBMIB with a *K_d_* value similar to other quinone binding proteins (25); it also forms a quinone adduct (26).

Figure 10 proposes a working model of signal perception mechanism of Hik2. In the absence of salt or redox stresses, the Hik2 is autokinase active and transfers phosphoryl groups to the response regulators Rre1 and RppA, while Phospho-Rre1 activates genes coding for phycobilisomes. However, in the presence of salt/osmotic stress, the active hexameric form of Hik2 is rearranged into the inactive tetrameric form. Rre1 and RppA therefore remain in their unphosphorylated states. As a result, Rre1 can no longer act as a repressor of salt/osmotic tolerance genes, in turn releasing the repression of their transcription. We are unable to determine the precise symmetry of the oligomeric forms described here. For example, the hexamers could be either trimers of dimers of 32-point group symmetry or hexamers of hexagonal symmetry (point group 6). Similarly, the tetramers could be dimers of dimers (point group 22), or tetramers of tetragonal symmetry (point group 4), and tetragonal tetramers may assemble into octamers as dimers of tetramers (point group 42). Conversion between hexamers and tetramers is likely to involve only oligomers of the aforementioned symmetries. This conversion between oligomers would be possible if conformations of the subunits change during the transition. The molecular basis of a mechanism by which elevated Na^+^ concentrations trigger recombination of hexamers into tetramers could therefore be twofold. Na^+^ may interfere with salt bridges that stabilise intersubunit interfaces at low salt concentrations. In addition, Na^+^ may induce conformational changes in the protein subunits thus leading to disruption of intersubunit interactions in the hexamers and favouring formation of new protein-protein interfaces that result in inactive tetramers.

Hik2 is of special interest because of the presence of its homolog in all known cyanobacteria as well as in chloroplasts (Table 1). In eukaryotes Hik2 has been identified as Chloroplast Sensor Kinase (CSK) that is encoded in the nucleus and synthesized in the cytosol for post-translational import and processing in chloroplasts (11, 27). CSK couples the redox state of the photosynthetic electron transport chain to chloroplast gene transcription by acting on plastid transcriptional regulators (25, 28). It will be important to determine whether the oligomerization that we propose as the basis of regulation by sodium ions is also a mechanism that extends to redox regulatory control of the activity of all Hik2 proteins, including CSK.

## MATERIALS AND METHODS

### Construction of recombinant plasmids

Coding sequences were cloned using the primer pairs listed in Table 3. These correspond to: the full-length *Synechocystis* sp. PCC6803 Hik2 (slr1147); and the *Thermosynechococcus elongatus* BP-1 (tlr0195) full-length, kinase domain corresponding to amino acid 142–386; and the DHp domain corresponding to amino acid 142–270. PCR products were digested with *NdeI* and *SalI* endonucleases (New England BioLabs) and cloned into pET-21b (Novagen) expression vector digested with *NdeI* and *XhoI*. The identities of the recombinant clones were confirmed by sequencing (results not shown).

**Table 3.**
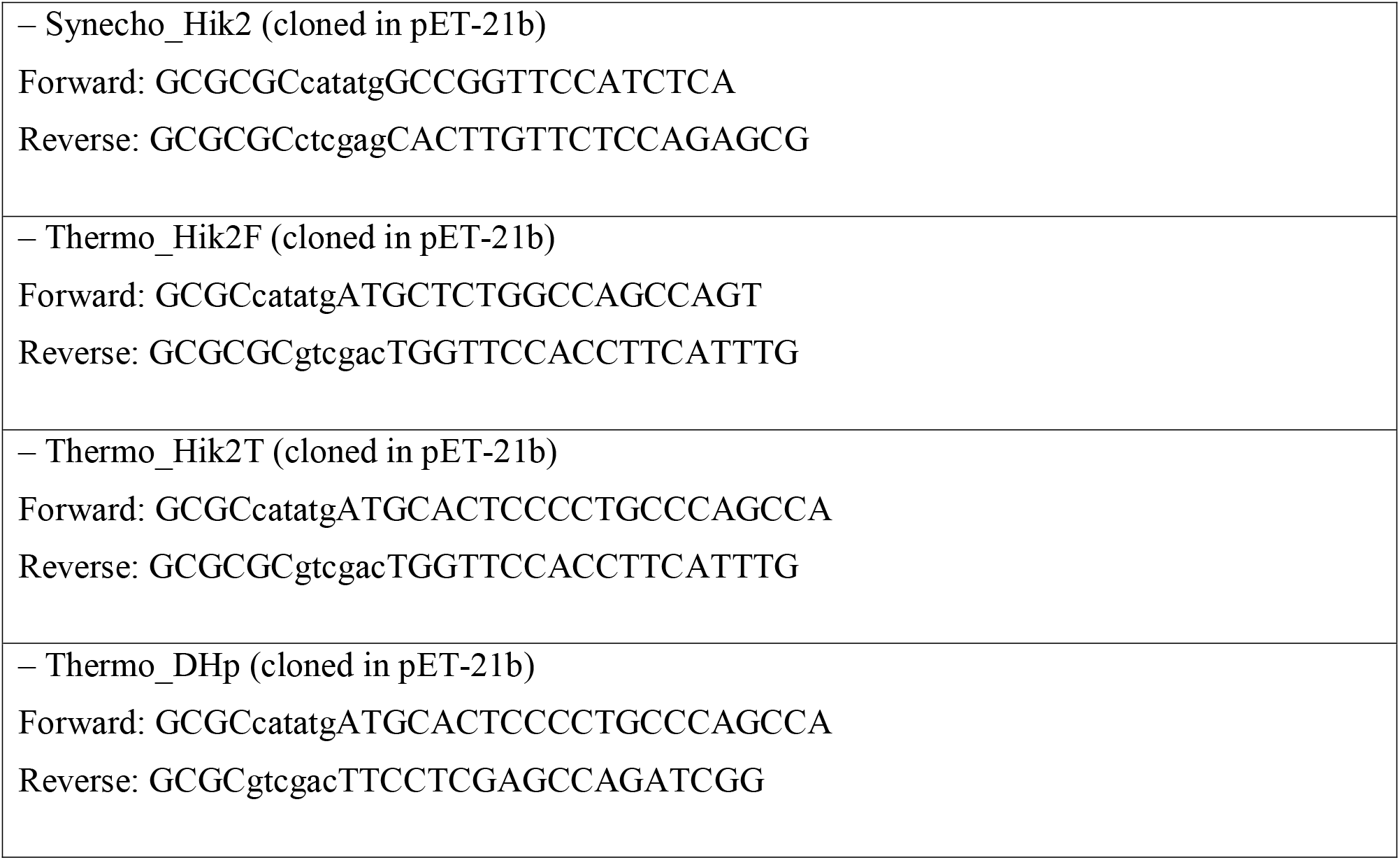
Primers used for Hik2 cloning. Sequences in lower case are restriction site overhangs. Sequences underlined are codons for alanine.

### Expression and purification of recombinant Hik2

Recombinant plasmids were transformed into *E. coli* BL21(DE3) chemically competent cells (Stratagene). Transformed bacterial colonies, grown on agar plates, were used to inoculate starter cultures (10 mL each) in Luria Broth (LB) growth media (29) with 100 *μ*g mL^−1^ ampicillin as the selectable marker. Each culture was grown overnight, diluted 1:100 in 1 L of LB media, and then grown at 37 °C to an optical density at 600 nm of ~ 0.55 before inducing protein expression with 0.5 mM IPTG (Melford). Bacterial cultures were then grown for a further 16 hours at 16 °C. Cells were harvested by centrifugation at 9,000 } g for 10 minutes at 4 °C. The pellet was suspended in a buffer containing 300 mM NaCl, 20 mM Tris-HCl adjusted to pH 7.4, 25 mM imidazole, and 1 mM PMSF, and the cells were then lysed with an EmulsiFlex-C3 homogenizer (Avestin). The lysate was separated by centrifugation at 39,000 } g for 20 min at 4 °C. The supernatant was applied to a Ni^2+^ affinity chromatography column (GE Healthcare) and purified according to the column manufacturer’s instructions.

### Chemical crosslinking

The full-length Hik2 protein was desalted into crosslinking reaction buffer (25 mM HEPES-NaOH at pH 7.5), 5 mM KCl, and 5 mM MgCl_2_) using a PD-10 desalting column (Amersham Biosciences). Chemical crosslinking was carried out in a total reaction volume of 20 *μ*L containing varying concentrations of Hik2 protein in crosslinking reaction buffer. The crosslinking agent dithiobis(succinimidylpropionate) (DSP) (30)was added from 24.73 mM stock solution in dimethyl sulfoxide to give a final DSP concentration of 2 mM. Reactions were incubated at 23 °C for 4 minutes. Reactions were stopped by addition of a solution containing 50 mM Tris-HCl and 10 mM glycine giving pH 7.5. The above reaction was repeated with 2 *μ*M Hik2 and varying concentrations of DSP. 2 *μ*g of crosslinked proteins were resolved upon 10 % SDS-PAGE (sodium dodecyl sulfate polyacrylamide gel electrophoresis) and the gel stained with Coomassie Brilliant Blue.

### In vitro autophosphorylation

Autophosphorylation was performed with 2 *μ*M of purified recombinant Hik2 protein in a kinase reaction buffer (50 mM Tris-HCl at pH 7.5, 50 mM KCl, 10 % glycerol, and 10 mM MgCl_2_) in a final reaction volume of 25 *μ*L. The autophosphorylation reaction was initiated by the addition of 5 *μ*L of a solution containing 2.5 mM disodium ATP (Sigma) with 2.5 *μ*Ci [γ-^32^P]-ATP (6000 Ci mmol^−1^) (PerkinElmer). Reactions were incubated for 15 seconds at 22 °C. Crosslinking was performed as above except that the autophosphorylation reaction was terminated, in this case, by addition of 6 *μ*L of 5-fold concentrated non-reducing Laemmli sample buffer (31). Reaction products were resolved using a 12 % nonreducing SDS-PAGE gel. The gel was rinsed with SDS running buffer and transferred into a polyethylene bag. The sealed bag was exposed to a phosphor plate overnight. The incorporated *γ*-^32^P was visualized using autoradiography.

### Sequence Analysis

Sequence similarity search was carried out with blastP and blastn (32) using the public databases Cyanobase (http://genome.kazusa.or.jp/cyanobase) and Joint Genome Institute (JGI) (http://www.jgi.doe.gov/). Domain prediction was carried out using the SMART database (http://smart.embl-heidelberg.de/smart/setmode.cgi?NORMAL=1)(33).

### Transmission electron microscopy (TEM) and single particle analyses

Five independent Hik2 samples were subjected in turn to a dilution series, applied to carbon-coated (thin-layer) copper 300-mesh EM grids (Agar, Ltd.), and negatively stained using freshly prepared 2 % uranyl acetate. A protein concentration that ensured an even spread of single particles over the carbon film surface was found for each sample and dilution. 201 micrographs (each being 2672 × 2672 pixels) were recorded using an Olympus Morada CCD camera system attached to a JEOL model 1230 TEM equipped with a tungsten filament and operating at 80,000 × magnification and 80 keV. This gave a sampling frequency of 5.962 Å per pixel at the specimen scale, however, a limitation of ~ 15 Å resolution is expected to result from the presence of the negative stain. The Fourier-space power spectrum was calculated for each micrograph and 41 micrographs were chosen for further single particle analysis on the basis that they displayed minimal drift and astigmatism and had a first minimum at better than ~ 17 Å resolution. Single particle complexes were floated out into boxes of 64 × 64 pixels in size. Given that no correction was applied for the contrast transfer function (CTF) the final class averages were low band-pass filtered to ~ 20 Å resolution. Initial single particle images were selected using the ‘boxer’ module of EMAN2 (34), with the boxing algorithm directed to pick automatically all possible single particles present, but not to band-pass or normalise. The Imagic-5 software environment (35) was then used for image normalisation, band-pass filtering, reference-free alignment and multi-variate statistical classification of the single particle image dataset.

## ACKNOWLEDGMENTS

IMI thanks Queen Mary University of London for a graduate teaching studentship. LW thanks the China Scholarship Council (CSC) and Queen Mary University of London for financial support. SP held a Leverhulme Trust early career postdoctoral research fellowship. JN is grateful for the continued support of the JST CREST Grant Number JPMJCR13M4, Japan. JFA acknowledges the support of research grant F/07 476/AQ and fellowship EM-2015-068 of the Leverhulme Trust.

## REFERENCES

1. Scheu PD, Liao YF, Bauer J, Kneuper H, Basche T, Unden G, Erker W. 2010. Oligomeric sensor kinase DcuS in the membrane of Escherichia coli and in proteoliposomes: chemical cross-linking and FRET spectroscopy. Journal of Bacteriology 192:3474–3483.

2. Swem LR, Kraft BJ, Swem DL, Setterdahl AT, Masuda S, Knaff DB, Zaleski JM, Bauer CE. 2003. Signal transduction by the global regulator RegB is mediated by a redox-active cysteine. EMBO Journal 22:4699–4708.

3. Filippou PS, Kasemian LD, Panagiotidis CA, Kyriakidis DA. 2008. Functional characterization of the histidine kinase of the E. coli two-component signal transduction system AtoS-AtoC. Biochimica et Biophysica Acta 1780:1023–1031.

4. Heermann R, Altendorf K, Jung K. 1998. The turgor sensor KdpD of Escherichia coli is a homodimer. Biochimica Et Biophysica Acta-Biomembranes 1415:114–124.

5. Cai SJ, Khorchid A, Ikura M, Inouye M. 2003. Probing catalytically essential domain orientation in histidine kinase EnvZ by targeted disulfide crosslinking. Journal of Molecular Biology 328:409–418.

6. Pan SQ, Charles T, Jin SG, Wu ZL, Nester EW. 1993. Preformed Dimeric State of the Sensor Protein Vira Is Involved in Plant-Agrobacterium Signal-Transduction. Proceedings of the National Academy of Sciences of the United States of America 90:9939–9943.

7. Surette MG, Levit M, Liu Y, Lukat G, Ninfa EG, Ninfa A, Stock JB. 1996. Dimerization is required for the activity of the protein histidine kinase CheA that mediates signal transduction in bacterial chemotaxis. Journal of Biological Chemistry 271:939–945.

8. Georgellis D, Kwon O, Lin ECC. 2001. Quinones as the redox signal for the Arc two-component system of bacteria. Science 292:2314–2316.

9. Malpica R, Franco B, Rodriguez C, Kwon O, Georgellis D. 2004. Identification of a quinone-sensitive redox switch in the ArcB sensor kinase. Proceedings of the National Academy of Sciences of the United States of America 101:13318–13323.

10. Ashby MK, Houmard J. 2006. Cyanobacterial two-component proteins: structure, diversity, distribution, and evolution. Microbiology and molecular biology reviews: MMBR 70:472–509.

11. Puthiyaveetil S, Kavanagh TA, Cain P, Sullivan JA, Newell CA, Gray JC, Robinson C, van der Giezen M, Rogers MB, Allen JF. 2008. The ancestral symbiont sensor kinase CSK links photosynthesis with gene expression in chloroplasts. Proceedings of the National Academy of Sciences of the United States of America 105:10061–10066.

12. Ibrahim IM, Puthiyaveetil S, Allen JF. 2016. A Two-Component Regulatory System in Transcriptional Control of Photosystem Stoichiometry: Redox-Dependent and Sodium Ion-Dependent Phosphoryl Transfer from Cyanobacterial Histidine Kinase Hik2 to Response Regulators Rrel and RppA. Frontiers in Plant Science 7.

13. Kobayashi I, Watanabe S, Kanesaki Y, Shimada T, Yoshikawa H, Tanaka K. 2017. Conserved two-component Hik34-Rre1 module directly activates heat-stress inducible transcription of major chaperone and other genes in Synechococcus elongatus PCC 7942. Molecular microbiology 104:260–277.

14. Ashby MK, Houmard J. 2006. Cyanobacterial two-component proteins: structure, diversity, distribution, and evolution. Microbiology and Molecular Biology Reviews 70:472–509.

15. Bilwes AM, Alex LA, Crane BR, Simon MI. 1999. Structure of CheA, a signal-transducing histidine kinase. Cell 96:131–141.

16. Marina A, Waldburger CD, Hendrickson WA. 2005. Structure of the entire cytoplasmic portion of a sensor histidine-kinase protein. EMBO J 24:4247–4259.

17. Surette MG, Levit M, Liu Y, Lukat G, Ninfa EG, Ninfa A, Stock JB. 1996. Dimerization is required for the activity of the protein histidine kinase CheA that mediates signal transduction in bacterial chemotaxis. Journal of Biological Chemical 271:939–945.

18. Wojnowska M, Yan J, Sivalingam GN, Cryar A, Gor J, Thalassinos K, Djordjevic S. 2013. Autophosphorylation Activity of a Soluble Hexameric Histidine Kinase Correlates with the Shift in Protein Conformational Equilibrium. Chemistry & Biology 20:1411–1420.

19. Gupta RS. 2009. Protein signatures (molecular synapomorphies) that are distinctive characteristics of the major cyanobacterial clades. International Journal of Systematic and Evolutionary Microbiology 59:2510–2526.

20. Mechaly AE, Sassoon N, Betton JM, Alzari PM. 2014. Segmental helical motions and dynamical asymmetry modulate histidine kinase autophosphorylation. PLoS Biol 12:e1001776.

21. Wang C, Sang J, Wang J, Su M, Downey JS, Wu Q, Wang S, Cai Y, Xu X, Wu J, Senadheera DB, Cvitkovitch DG, Chen L, Goodman SD, Han A. 2013. Mechanistic insights revealed by the crystal structure of a histidine kinase with signal transducer and sensor domains. PLoS Biol 11:e1001493.

22. Bhate MP, Molnar KS, Goulian M, DeGrado WF. 2015. Signal transduction in histidine kinases: insights from new structures. Structure 23:981–994.

23. Wang LC, Morgan LK, Godakumbura P, Kenney LJ, Anand GS. 2012. The inner membrane histidine kinase EnvZ senses osmolality via helix-coil transitions in the cytoplasm. The EMBO journal 31:2648–2659.

24. Allen JF. 1993. Redox control of transcription: sensors, response regulators, activators and repressors. FEBS Letter 332:203–207.

25. Puthiyaveetil S, Ibrahim IM, Allen JF. 2013. Evolutionary rewiring: a modified prokaryotic gene regulatory pathway in chloroplasts. Philosophical Transactions of the Royal Society B: Biological 367.

26. Ibrahim IM, Puthiyaveetil S, Khan C, Allen JF. 2016. Probing the nucleotide-binding activity of a redox sensor: two-component regulatory control in chloroplasts. Photosynth Research doi:10.1007/s11120-016-0229-y.

27. Puthiyaveetil S, Ibrahim IM, Allen JF. 2012. Oxidation-reduction signalling components in regulatory pathways of state transitions and photosystem stoichiometry adjustment in chloroplasts. Plant Cell and Environment 35:347–359.

28. Puthiyaveetil S, Ibrahim IM, Jelicic B, Tomasic A, Fulgosi H, Allen JF. 2010. Transcriptional control of photosynthesis genes: the evolutionarily conserved regulatory mechanism in plastid genome function. Genome Biology Evolution 2:888–896.

29. Sambrook J, Fritsch EF, Maniatis T. 1989. Molecular cloning a laboratory manual second edition.

30. Lomant AJ, Fairbanks G. 1976. Chemical probes of extended biological structures: synthesis and properties of the cleavable protein cross-linking reagent [35S]dithiobis(succinimidyl propionate). Journal of Molecular Biology 104:243–261.

31. Laemmli UK. 1970. Cleavage of structural proteins during assembly of head of bacteriophage-T4. Nature 227:680.

32. Altschul SF, Gish W, Miller W, Myers EW, Lipman DJ. 1990. Basic local alignment search tool. Journal of Molecular Biology 215:403–410.

33. Schultz J, Milpetz F, Bork P, Ponting CP. 1998. SMART, a simple modular architecture research tool: identification of signaling domains. Proceedings of the National Academy of Sciences of the United States of America 95:5857–5864.

34. Tang G, Peng L, Baldwin PR, Mann DS, Jiang W, Rees I, Ludtke SJ. 2007. EMAN2: an extensible image processing suite for electron microscopy. J Struct Biol 157:38–46.

35. van Heel M, Harauz G, Orlova EV, Schmidt R, Schatz M. 1996. A new generation of the IMAGIC image processing system. J Struct Biol 116:17–24.

